# Dynamic Analysis of Alternative Polyadenylation from Single-Cell RNA-Seq (scDaPars) Reveals Cell Subpopulations Invisible to Gene Expression Analysis

**DOI:** 10.1101/2020.09.23.310649

**Authors:** Yipeng Gao, Lei Li, Christopher I. Amos, Wei Li

## Abstract

Alternative polyadenylation (APA) is a major mechanism of post-transcriptional regulation in various cellular processes including cell proliferation and differentiation, but the APA heterogeneity among single cells remains largely unknown. Single-cell RNA sequencing (scRNA-seq) has been extensively used to define cell subpopulations at the transcription level. Yet, most scRNA-seq data have not been analyzed in an “APA-aware” manner. Here, we introduce scDaPars, a bioinformatics algorithm to accurately quantify APA events at both single-cell and single-gene resolution using standard scRNA-seq data. Validations in both real and simulated data indicate that scDaPars can robustly recover missing APA events caused by the low amounts of mRNA sequenced in single cells. When applied to cancer and human endoderm differentiation data, scDaPars not only revealed cell-type-specific APA regulation but also identified cell subpopulations that are otherwise invisible to conventional gene expression analysis. Thus, scDaPars will enable us to understand cellular heterogeneity at the post-transcriptional APA level.

## Introduction

Alternative polyadenylation (APA) is a major mechanism of post-transcriptional regulation under diverse physiological and pathological conditions (Elkon et al. 2013; Tian and Manley 2017). The process of polyadenylation involves endonucleolytic cleavage of the nascent RNA followed by synthesis of a poly(A) tail on the 3’ terminus (Tian and Manley 2017). By using different polyadenylation sites (poly(A) sites), which are defined by flanking RNA sequence motifs, APA can generate mRNA isoforms with various 3’-untranslated regions (3’UTRs) in the majority of human genes (Derti et al. 2012; Tian and Manley 2017). While APA in most cases does not alter the protein-coding regions in those mRNA isoforms, it disrupts important *cis*-regulatory elements located in the 3’UTRs, including adenylate-uridylate-rich elements (ARE) and binding sites of miRNAs and RNA binding proteins, resulting in altered mRNA stability, localization and translation efficiency (Garneau et al. 2007; An et al. 2008; Hoffman et al. 2016).

Next-generation sequencing technologies have revolutionized our understanding of APA over the last decade, illustrating both the pervasiveness of dynamic APA events and complexity of the APA regulatory processes. Recently, multiple studies have shed light on the global regulation of APA in response to changes in cell proliferation and cell differentiation in human diseases including cancer (Tian and Manley 2017; Gruber and Zavolan 2019). Both proliferating cells and transformed cells often express a multitude of alternative mRNA isoforms with shortened 3’UTRs through APA (Sandberg et al. 2008), leading to the activation of several proto-oncogenes such as cyclinD1, by escaping miRNA-mediated repression (Mayr and Bartel 2009). On the other hand, 3’UTR lengthening is more prevalent in cell differentiation (Ji et al. 2009; Ji and Tian 2009). For example, progressive 3’UTR lengthening is observed during mouse embryonic development (Ji et al. 2009), and the generation of induced pluripotent stem cells (iPSCs) (dedifferentiation) is accompanied by global 3’UTR shortening (Ji and Tian 2009). Besides regulating cognate transcripts in *cis*, APA-induced 3’UTR changes can also disrupt competing endogenous RNA (ceRNA) regulation in *trans*, thus repressing several crucial tumor suppressors such as PTEN in breast cancer (Park et al. 2018). Although these observations imply a possible cell-state- or cell-type-dependent manner of APA regulation, the variability of APA among individual cells and the utility of APA in revealing novel cell subpopulations remains largely unknown.

Single-cell RNA sequencing (scRNA-seq) has become one of the most widely used technologies in biomedical research by providing an unprecedented opportunity to quantify the abundance of diverse transcript isoforms among individual cells (Shapiro et al. 2013; Saliba et al. 2014). However, methods to quantify relative APA dynamics across single cells remain underdeveloped. Recently, Velten et al. (Velten et al. 2015) developed an experimental protocol BATseq to quantify various 3’UTR-isoforms at the single-cell resolution. By integrating the standard scRNA-seq protocol and the 3’ enriched bulk RNA-seq protocol, Velten et al. found that cell types can be well separated based exclusively on their 3’UTR isoform usages, indicating that APA is a molecular feature intrinsic to cell states (Velten et al. 2015). While a compelling method, BATseq is hampered by its low sensitivity (~5%) and high procedural complexity (Chen et al. 2017), thereby not being widely adopted in practice. In contrast, standard scRNA-seq data is widely available, yet most of the scRNA-seq data has not been analyzed in an “APA-aware” manner. Since scRNA-seq only captures a small fraction (typically 5%-15%) of the total mRNAs in each cell (Stegle et al. 2015), it can falsely quantify genes, especially lowly expressed ones, as unexpressed; this phenomenon is termed as “dropout”. Existing bulk RNA-seq based APA methods such as DaPars (Xia et al. 2014) cannot overcome this vexing challenge when applied directly to scRNA-seq data, as they would lead to a high degree of sparsity in the resulting APA profiles. To address this sparsity, recently published computational approaches such as scDAPA (Ye et al. 2020) scAPA (Shulman and Elkon 2019) extract and combine reads from cells aggregated based on pre-defined cell types. Alternatively, another study (Kim et al. 2019) aggregates individual genes into “meta-genes” with reference to common functionality. While these strategies cope with the problem of sparsity to some extent, they fail to retain the single-cell or single-gene resolution (Supplementary table 1).

To fill this knowledge gap, we developed scDaPars (**D**ynamic analysis of **A**lternative **P**oly**A**denylation from **scR**NA-**S**eq), a bioinformatics algorithm for quantifying and recovering APA usage at the single-cell and single-gene resolution using standard scRNA-seq data. Since APA is reported to be regulated in a cell-state- or cell-type-specific manner, scDaPars employs a regression model that enables sharing of APA information across related cells to tackle the sparsity. The regression model couples the tasks of borrowing APA information of the same gene in neighboring cells and of transferring APA information from genes unlikely to be affected by dropout events (robust genes) to dropout genes, achieving considerable robustness when applied to noisy scRNA-seq data. To the best of our knowledge, scDaPars is the first single-cell- and single-gene-level APA quantification method for analyzing standard scRNA-seq data.

## Results

### Overview of the scDaPars algorithm

Figure 1 presents a schematic illustration of the scDaPars algorithm (see “Methods” for detailed definition and computational procedures). Given a scRNA-seq dataset, scDaPars first calculates raw relative APA usage, measured by the percentage of distal poly(A) site usage index (PDUI), based on the two-Poly(A)-site model introduced in DaPars (Xia et al. 2014). scDaPars takes scRNA-seq genome coverage data as input and forms a linear regression model to jointly infer the exact location of proximal poly(A) sites by minimizing the deviation between the observed read density and the expected read density in all single cells. The relative APA usage is then quantified as the proportion of the estimated abundances of transcripts with distal poly(A) sites (longer 3’UTRs) out of all transcripts (longer and shorter 3’UTRs), and therefore, genes favoring distal poly(A) site usage (long 3’ UTRs) will have PDUI values near 1, whereas genes favoring proximal poly(A) site usage (short 3’ UTRs) will have PDUI values near 0. This step (step (I)) will generate a PDUI matrix with rows representing genes and columns representing single cells. Of note, the raw PDUI values can only be estimated for genes with sufficient read coverages (default coverage of 5 reads per base), which automatically separates genes into robust genes (genes unaffected by dropout events) and dropout genes for further analysis. Due to the intrinsically low coverage of scRNA-seq data (Brennecke et al. 2013), the resulting PDUI matrix from step (I) is overly sparse with widespread missing data. To further recover the complete PDUI matrix, we develop a new imputation method by sharing APA information across different cells. For a given cell, scDaPars begins by constructing a nearest neighbor graph based on the sparse PDUI matrix generated in step (I) (Fig. 1) to identify a pool of candidate neighboring cells that have similar APA profiles (step (II)). Finally, scDaPars uses a non-negative least square (NNLS) regression model to refine neighboring cells based on robust genes and then borrow APA information in these neighboring cells to impute PDUIs of dropout genes in each cell (step (III)).

**Figure 1.**
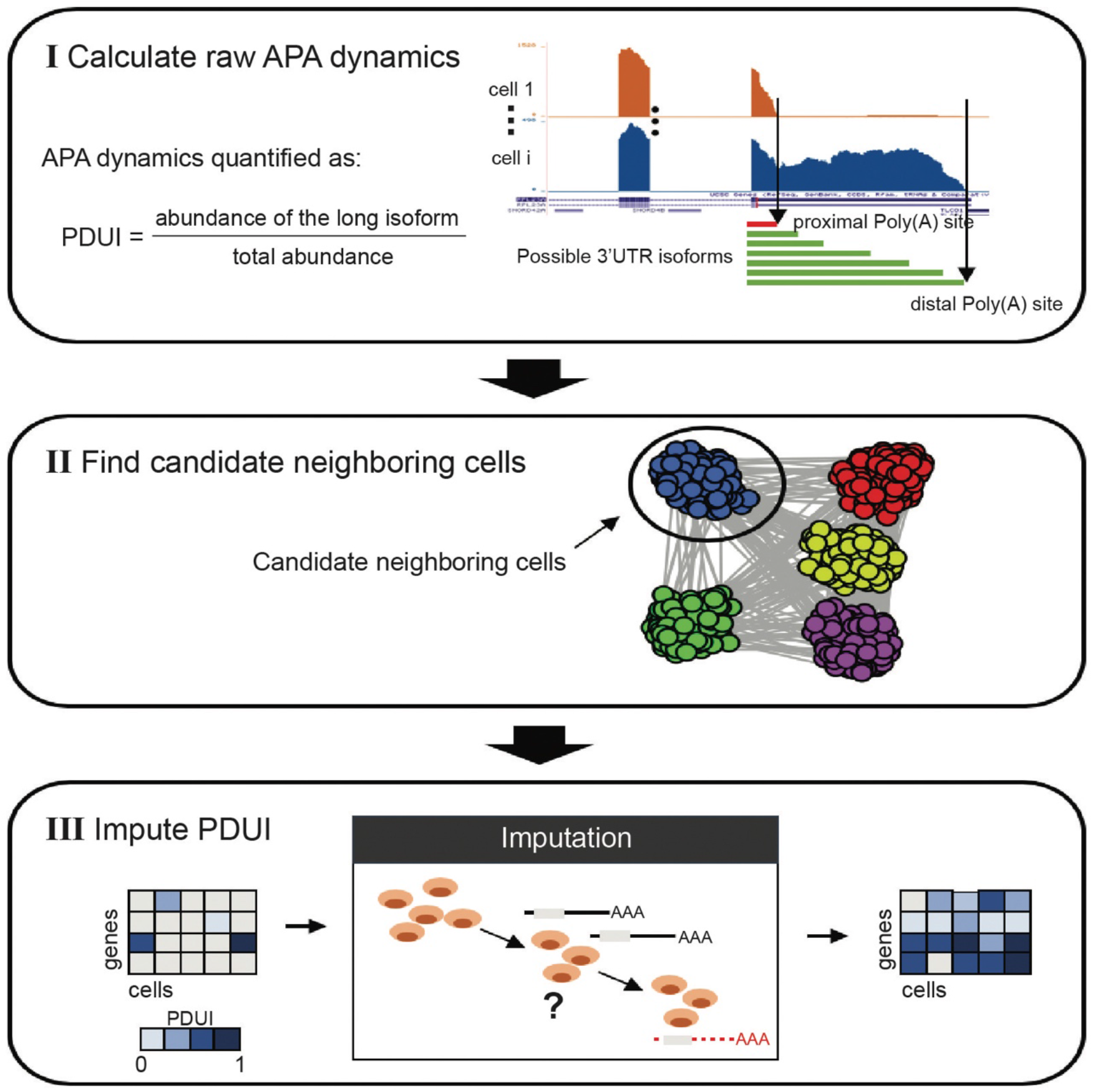
A schematic illustration of the scDaPars algorithm. (I) scDaPars predicts both distal and proximal poly(A) sites by joint analysis of all single-cell samples and quantifies the raw relative APA usage by the proportion of estimated abundances of transcripts with distal poly(A) sites (long isoform). (II) scDaPars determines potential neighboring cells by applying community detection methods in APA profiles generated in step(I). (III) scDaPars uses NNLS regression model to refine neighboring cells and impute missing values by borrowing APA information from neighboring cells.

### Evaluation of the Accuracy and Robustness of scDaPars

To quantitatively evaluate the accuracy of imputed APA events by scDaPars, we used 384 scRNA-seq libraries of individual human peripheral blood cells (PBMCs) sequenced by Smart-seq2 (Picelli et al. 2013) protocol and a matched bulk RNA-seq library from a benchmark study by Ding et al. (Ding et al. 2020). Since we can estimate poly(A) sites and quantify differential poly(A) sites usages with high sensitivity and specificity in bulk RNA-seq datasets (Xia et al. 2014), we treated the results from the matched bulk sample as pseudo-gold standard for the following evaluation.

First, we showed that scDaPars reliably identified the location of proximal poly(A) sites in single cells. We found that ~84% of poly(A) sites predicted from scRNA-seq data are within 100bp of those predicted in bulk, whereas only ~44% of randomly selected sites from 3’UTR regions are within 100bp of bulk predictions (Fig. 2a). Reassuringly, ~66.2% of poly(A) sites predicted from scRNA-seq data also overlapped with annotated poly(A) sites complied from RefSeq, ENSEMBL, UCSC gene models and PolyA_DB (Wang et al. 2017) within 100bp, and this overlap showed an approximately fivefold enrichment compared with random sites (Fig. 2b). In addition, canonical PolyA signal (PAS) AATAAA was successfully identified by *de novo* motif analysis(Bailey 2011) within the upstream (−100bp) sequence of single-cell predicted poly(A) sites with a p-value (P = 1.2e-44) similar to that of bulk samples (P = 5.4e-48) (Fig. 2c, supplementary Fig.1), supporting the validity of scDaPars’s prediction of poly(A) sites.

**Figure 2.**
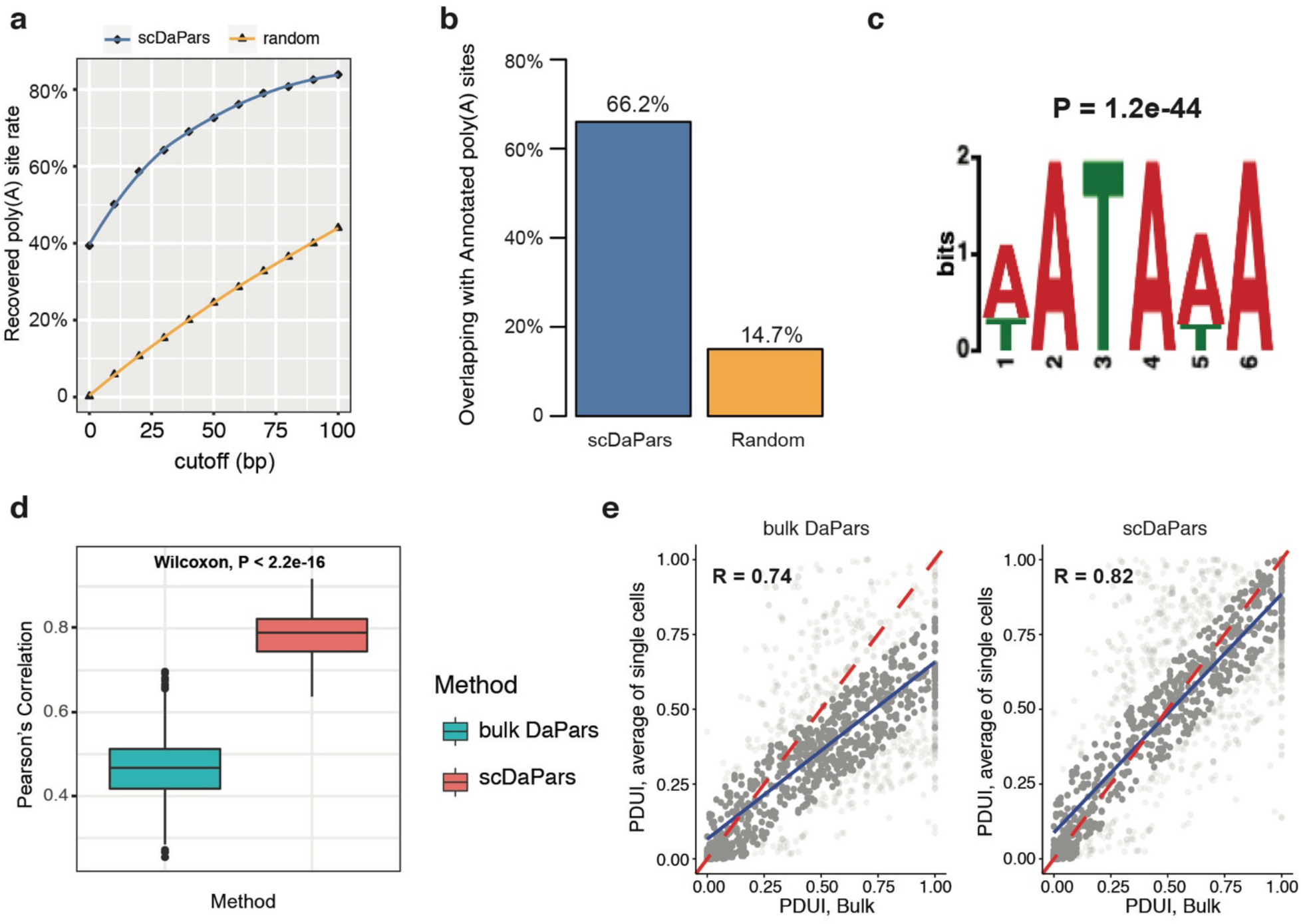
Evaluation of APA detection accuracy of scDaPars using human PBMCs datasets. (a) Fraction of recovered bulk-predicted poly(A) sites using scDaPars or random control. Poly(A) sites predicted in scRNA-seq are considered true if they are located within cutoff distance from the bulk results. The cutoffs range from 0 to 100bp with 10bp increment. (b) Percentage of scDaPars predicted poly(A) sites or random control overlapped with annotated poly(A) sites from RefSeq, ENSEMBL, UCSC gene models and PolyA_DB. (c) The top-scoring signal identified by de novo motif analysis (DREME) from the upstream (−100bp) of scDaPars predicted poly(A) sites from single cells. (d) Boxplot showing Pearson’s correlations between PDUI values of B-cell pairs estimated by bulk DaPars and scDaPars (Wilcoxon test P < 2.2e-16). (e) Scatter plots of PDUI values between average of all single cells and bulk results estimated by bulk DaPars (left) and scDaPars (right). Red line represents the theoretical linear relationships between bulk and single cell PDUIs, and blue represents the actual linear relationships estimated from data.

Next, we showed that scDaPars was able to recover APA usage for genes affected by dropouts in scRNA-seq data. APA is found to be uniquely regulated in distinct immune cell types in PBMCs (Kim et al. 2019). Yet the median Pearson’s correlation between APA (PDUI values) of single-cell pairs in the same B cell cluster is only 0.46 when PDUI values were calculated by DaPars (our previous method for bulk RNA-seq) due to dropout effects (Fig. 2d). In contrast, scDaPars successfully recovered PDUI values for most of the affected dropout genes (Supplementary Fig. 2) and increased the median cell-cell correlation by a large margin (0.79) (P < 2.2e-16) (Fig. 2d). We further compared the average APA usage of all single cells with the bulk results. The Pearson’s correlation between the average PDUI values of single cells and those of the bulk increased from 0.74 to 0.82 after scDaPars imputation (Fig. 2e). Notably, even though the correlation increase was not large, the regression slope increased significantly from 0.59 (DaPars) to 0.8 (scDaPars) (P = 4.89e-26), indicating APA usages quantified by scDaPars better represents the linear relationship between the average of single-cell APA dynamics and the corresponding bulk APA dynamics.

Finally, we used a simulation study to illustrate scDaPars’s ability to identify dynamic APA events between two cell types. We created a synthetic PDUI matrix of Naïve and activated CD4 T cells based on bulk RNA-seq data from the DICE project (Schmiedel et al. 2018) (see “Methods”). The Naïve and activated CD4 T cells are clearly distinguishable using the reference APA profiles estimated from bulk samples (Fig. 3a).

**Figure 3.**
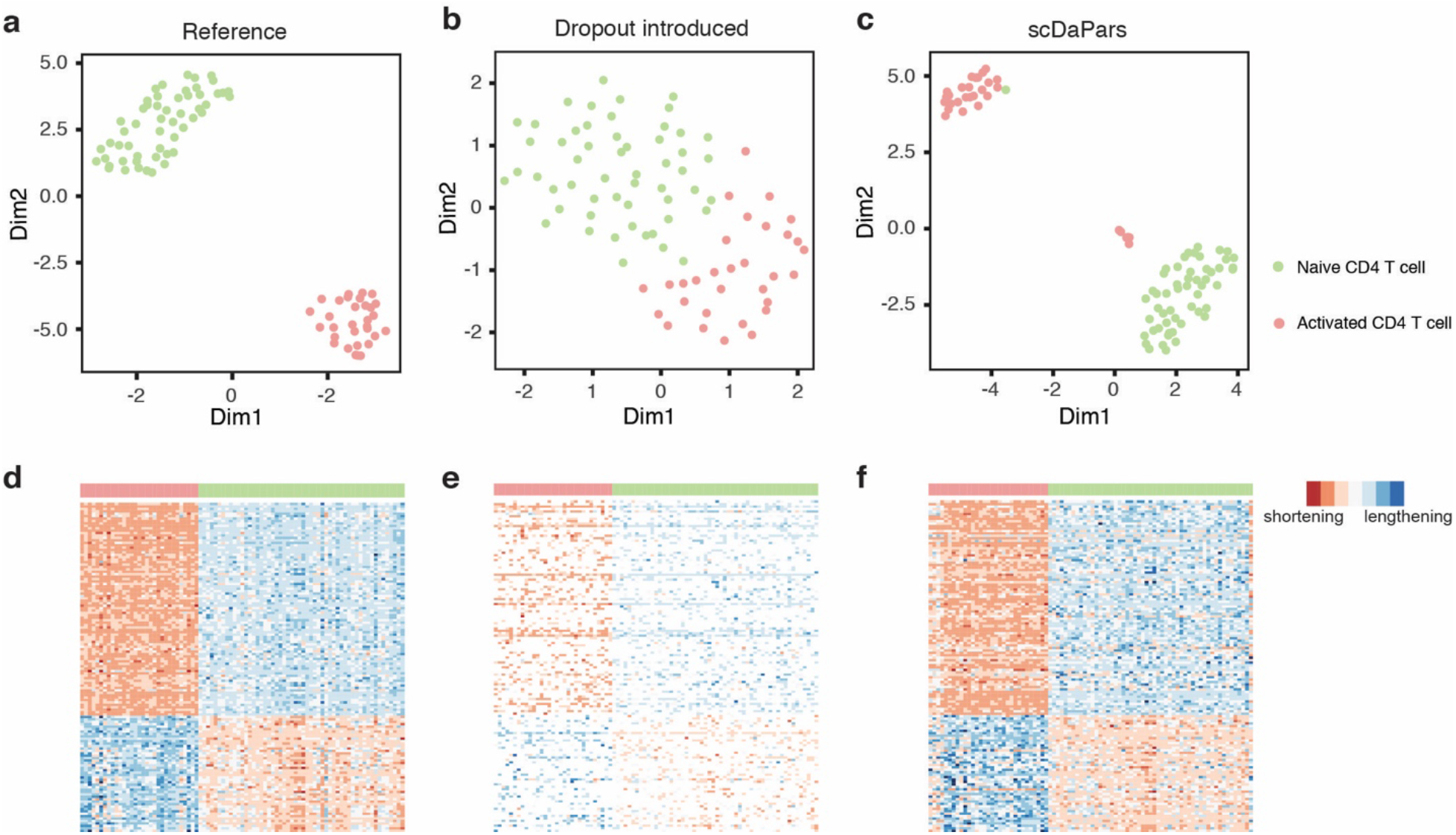
Evaluation of scDaPars in identifying dynamic APA events between two cell types using Naïve and activated CD4 T cells. (a) – (c) Scatterplots showing UMAP results of 54 Naïve CD4 T cells and 31 activated CD4 T cells based on (a) Reference APA profiles or (b) Dropout events introduced APA profiles or (c) scDaPars corrected APA profiles. (d) – (f) Heatmaps showing APA profiles of 136 differential APA genes (PDUI differences >= 0.2) in the (d) reference data (e) dropout event introduced data and (f) scDaPars corrected data. Rows represent differential APA genes and columns represent cells. 88 out of 136 differential APA genes have shorter 3’UTRs in activated CD4 T cells.

Additionally, the reference data showed a strong inclination of 3’UTR shortening in activated CD4 T cells (P = 3.8e-4) (Fig. 3d), in line with previous reports that 3’UTR shortening is widely observed upon activation of T cells (Sandberg et al. 2008). However, manually introduced dropout events obscured this differential 3’UTR pattern, in which only ~38% of differential APA genes remained, and the two cell types became less separated by their APA profiles (Figs. 3b and 3e). After we applied the imputation steps of scDaPars, ~79% of differential APA genes are recovered and the clear separation of these two cell types was restored (Figs. 3c and 3f). We further examined the robustness of scDaPars against varying dropout rates. Even though the accuracy of dynamic APA events identified by scDaPars decreased as the dropout rate increased, scDaPars could still achieve > 0.75 area under the receiver operating characteristics (ROC) curve when the proportion of dropout events was as high as 70% (Supplementary Fig. 3).

### scDaPars outperforms existing methods by providing single-cell-resolution APA quantification applicable to both 3’ end and full-length scRNA-seq data

Several bioinformatics tools have been developed to analyze APA dynamics using scRNA-seq data (i.e. scDAPA (Ye et al. 2020) and scAPA (Shulman and Elkon 2019)), yet, unlike scDaPars, they were not designed to quantify APA dynamics at the singlecell resolution. In addition, during the preparation of this manuscript, we noticed another method Sierra (Patrick et al. 2020), which detects differential transcript usage in scRNA-seq data, may also be used for quantifying dynamic APA events. To illustrate the superiority of scDaPars over these existing methods, we applied scDaPars, scAPA and Sierra to a benchmark 10X Chromium dataset containing 3362 PBMCs (Ding et al. 2020) (see “Methods”). scDAPA was excluded from this study since it identifies APA events by pair-wise comparison without quantifying single-cell APA usage. The APA usage quantified by scDaPars generated clear and compact immune cell clusters (Fig. 4a). In contrast, although Sierra outperformed scAPA and was able to identify B cell and CD14+ monocytes, both Sierra and scAPA failed to distinguish the five immune cell types (Fig. 4b, c). We also used silhouette analysis to quantitatively assess the resulting clusters. Compared with scAPA and Sierra, scDaPars showed higher silhouette coefficients which indicated the clustering results from scDaPars are more congruent with the true biological cell types (Fig. 4d, e and f). More importantly, both scAPA and Sierra rely on peak calling using 3’ end enriched reads in 10X Chromium to quantify APA usage and thus are not applicable to data generated by full-length sequencing protocols like Smart-Seq2 which do not contain enriched peaks in the 3’UTR regions (Picelli et al. 2013).

**Figure 4.**
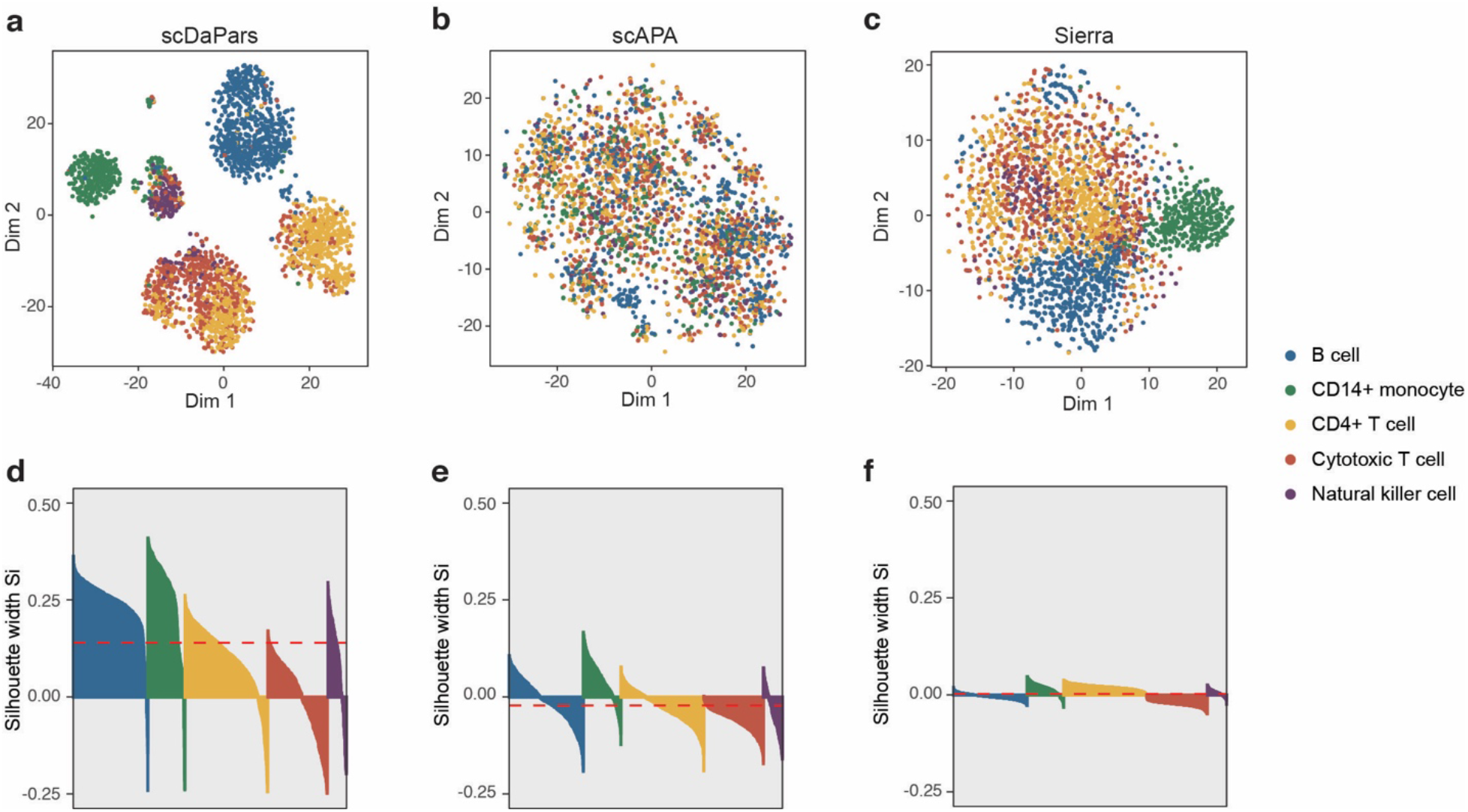
scDaPars outperforms existing methods by quantifying APA dynamics in single-cell resolution. (a) – (c) Scatterplots showing UMAP results of 3362 PBMCs based on (a) scDaPars quantified APA usage or (b) scAPA quantified APA usage or (c) Sierra quantified APA usage. (d) – (f) Silhouette plots for clustering results from (d) scDaPars, (e) scAPA and (f) Sierra. The x-axis represents cells and y-axis is the corresponding silhouette measure Si for each cell. The silhouette coefficient measures how similar a cell is to its own cluster compared to other clusters; therefore, a higher silhouette coefficient indicates better clustering results and a negative coefficient may suggest the cell is assigned to the wrong cluster. The red dashed line is the average Si for all cells.

### scDaPars revealed intrinsic tumor APA variations and immune cell subpopulations in primary breast cancer

Global-scale coordinated APA events are commonly observed in cancers (Xia et al. 2014), and APA induced 3’UTR shortening was shown to be associated with tumor aggressiveness and poor survival of cancer patients (Lembo et al. 2012; Xia et al. 2014). However, knowledge of APA dynamics in cancer has been largely derived from bulk RNA-seq studies. Therefore, while global APA variations between tumor and normal cells have been well characterized, little is known about the intertumoral APA heterogeneity at the single-cell resolution. To illustrate scDaPars’ capacity of characterizing single-cell APA variations in cancers, we applied scDaPars to a Smart-seq2 (Picelli et al. 2013) scRNA-seq dataset containing 563 single cells from 11 breast cancer patients (Chung et al. 2017). In consistent with bulk results, 3’UTRs were significantly shortened in tumor cells compared to normal cells (P < 2.2e-16) (Fig. 5a). Interestingly, even PDUI values before scDaPars imputation could separate tumor cells from non-tumor cells with effectiveness comparable to that of gene expression values (UMAP (McInnes et al. 2018) visualization in supplementary Fig. 4a), suggesting an important role of dynamic APA events in breast cancer progression. As expected, scDaPars imputed APA profiles showed a much better separation between tumor and non-tumor groups (Fig. 5b).

**Figure 5.**
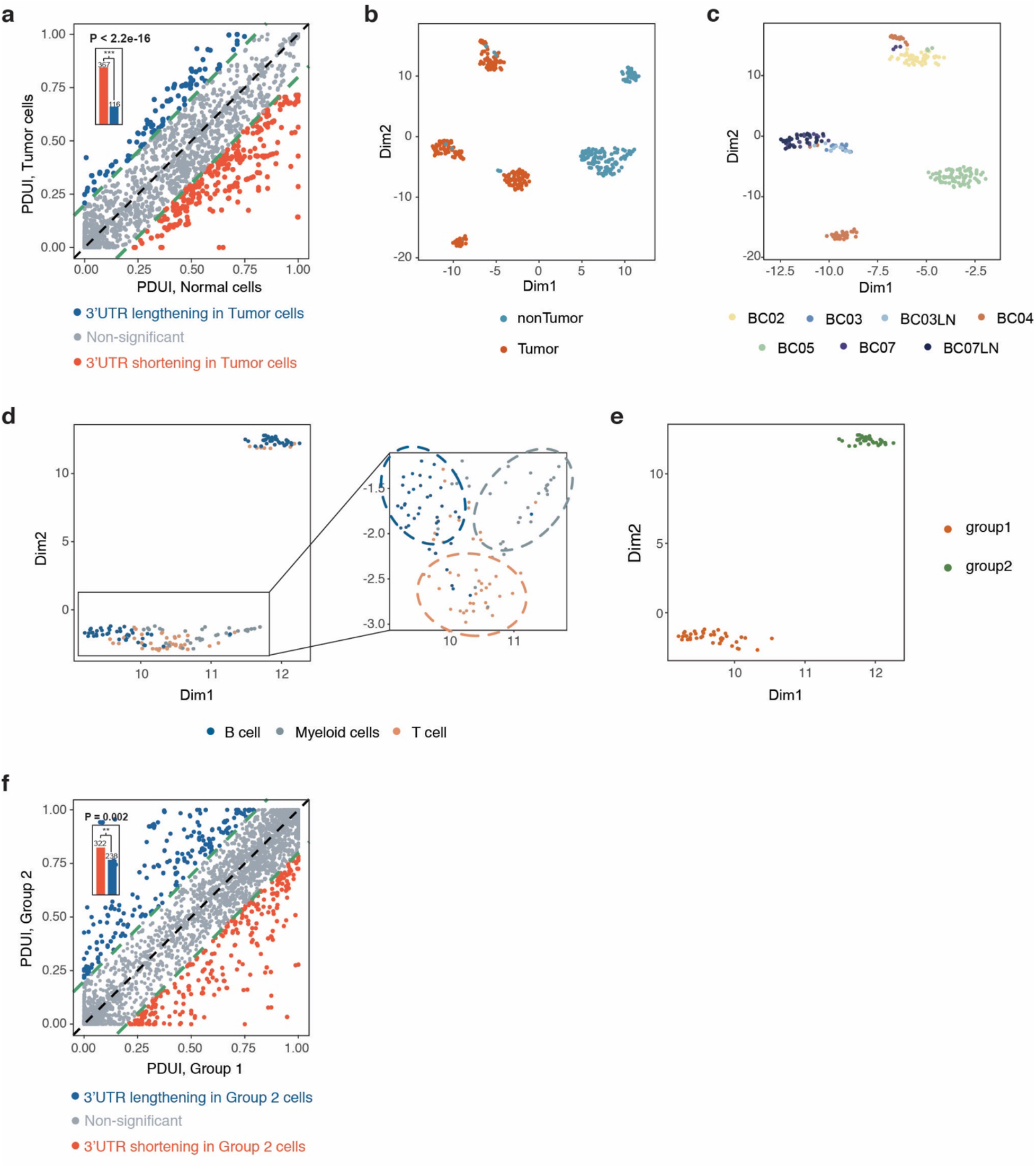
scDaPars reveals tumor-specific and immune-cell-type specific APA dynamic in primary breast cancer. (a) Scatter plot of PDUI values in Tumor and Normal cells. For each gene, the mean PDUI values in tumor cells (y-axis) are plotted against that in normal cells (x-axis). Genes with significant shortening or lengthening 3’UTR (PDUI difference >= 0.2) in tumor cells are shown in red and blue. Bar plot shows the number of genes with significant shortening or lengthening in tumor cells and p-value is calculated using single-tailed binomial test. (b) Scatter plot gives UMAP results calculated from scDaPars restored APA profiles. Each dot represents a cell, and cells are labeled based on cell index provided in the original publication. (c) Scatter plot of UMAP results of tumor cells. Cells are labeled by patient information. (d) Scatter plot of UMAP results of immune cells. Cells are labeled by cell type information. (e) Scatter plot of UMAP results of B cells based on scDaPars results. (f) Scatter plot of PDUI values in group 1 B cell and group 2 B cell. For each gene, the mean PDUI values in group 2 B cells (y-axis) are plotted against that in group 1 B cells (x-axis). Genes with significant shortening or lengthening 3’UTR (PDUI difference >= 0.2) in group 2 B cells are shown in red and blue. Bar plot shows the number of genes with significant shortening or lengthening in group 2 cells.

To further elucidate APA variations among cell subgroups, we analyzed APA profiles of tumor and non-tumor cells separately. On the one hand, contrary to a previous single-cell APA analysis performed on aggregated “meta-genes” in the same breast cancer dataset (Chung et al. 2017), which showed that no differences in APA were associated with cancer subtypes or patients (Kim et al. 2019), we found that tumor cells were separated into patient-specific clusters based on scDaPars-imputed APA profiles (Fig. 5c), showing evidence of intertumoral APA heterogeneity as well as scDaPars’s advantage over existing method. On the other hand, non-tumor cells, which were derived from the same group of patients as tumor cells, were clustered mainly according to their cell types (B cells, Myeloid cells and T cells) instead of patients (Fig. 5d, Supplementary Fig. 4b). This result not only reaffirmed that dynamic APA events are cell-type specific characteristics of immune cells, but also indicated that the patient-specific APA profiles observed in tumor cells were unlikely due to batch effects in patient samples but rather reflected true intertumoral variations in APA.

In addition, we observed that B cells were further classified into two cell subgroups based on scDaPars-imputed APA profiles (Fig. 5e) with group 2 B cells showed global 3’UTR shortening compared with group 1 B cells (P = 0.002) (Fig. 5f). Interestingly, we found that most B cell proliferation signature genes from the literature (Chung et al. 2017) were significantly more upregulated in group 2 B cells compared to group 1 B cells (Supplementary Fig. 5, Supplementary Table 2), suggesting that group 2 B cells may represent proliferating B cells. Indeed, the proliferating and non-proliferating cells determined by the expression of B cell proliferating marker genes are highly congruent with scDaPars derived cell subgroups (Supplementary Figs. 6a, b). These results are also consistent with previous reports that proliferating cells (i.e. group 2 cells) express more isoforms with shortened 3’UTRs through APA (Sandberg et al. 2008). Surprisingly, however, expression analysis of all genes failed to identify these B cell subgroups (Supplementary Fig. 6c), revealing the potential benefits of APA analysis in delineating cell subpopulations. In summary, scDaPars improves the characterization of APA variations and cell subpopulations in single cells.

### scDaPars enables identification of novel cell subpopulations invisible to conventional gene expression analysis in endoderm differentiation

As APA patterns appear to be globally regulated in cell differentiation (Ji et al. 2009; Tian and Manley 2017) (i.e. decreased proximal poly(A) sites usage in more differentiated states of embryonic development), we hypothesized that they could provide a new aspect to identify cell subpopulations during differentiation. To test this hypothesis, we applied scDaPars to a time-course Smart-seq2 (Picelli et al. 2013) scRNA-seq dataset containing 758 cells sequenced at 0, 12, 24, 36, 72 and 96 h of differentiation during human definitive endoderm (DE) emergence (Chu et al. 2016). scDaPars revealed clear and compact cell clusters from each time point along the differentiation process (Fig. 6a). Interestingly, dimension 2 of the UMAP projection of raw PDUI values reconstructed single-cell orders matching the true differentiation time points, reflecting the global APA dynamics during cell differentiation (Supplementary Fig. 7).

**Figure 6.**
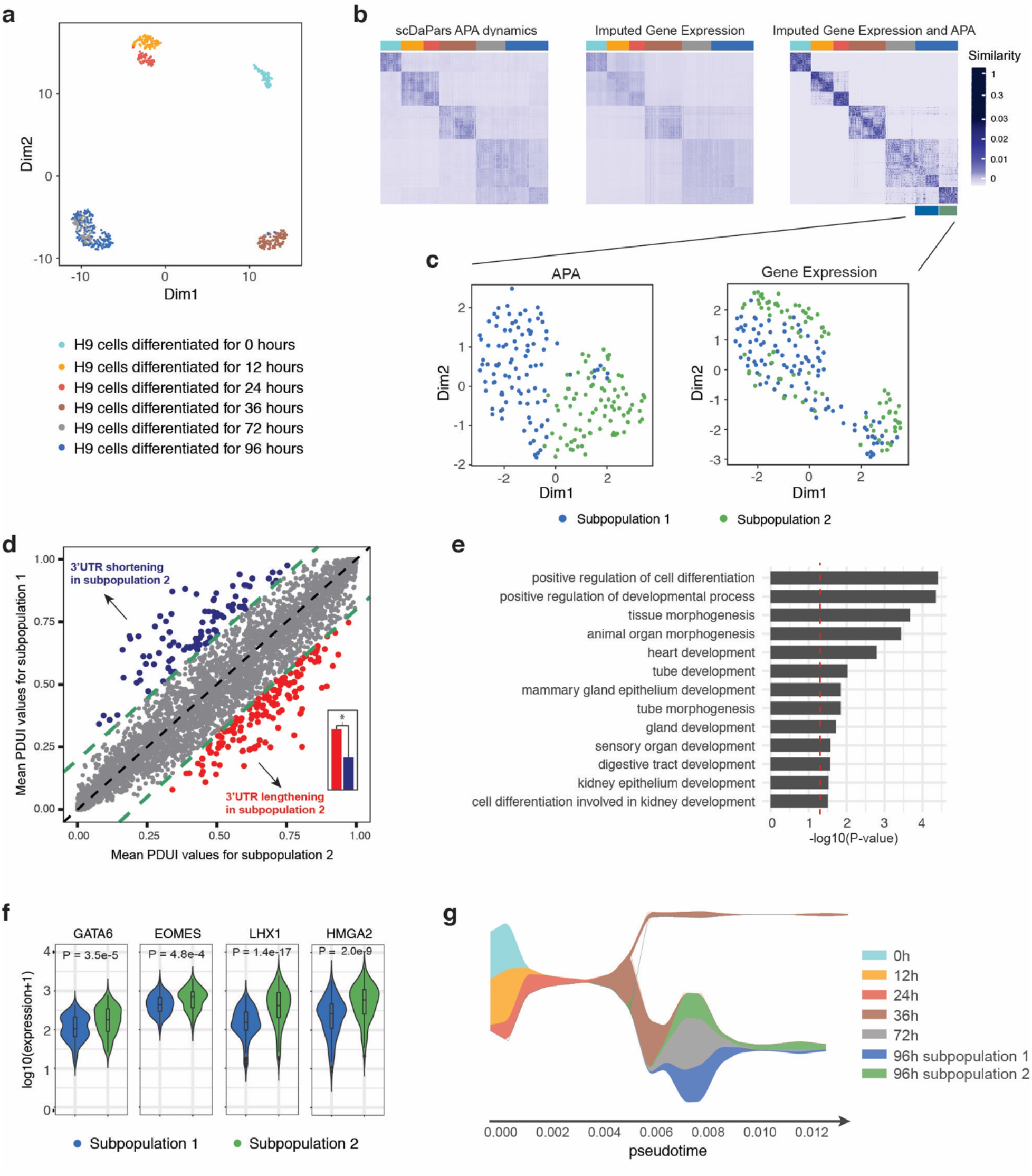
scDaPars helps identify novel cell subpopulations during human embryonic development. (a) Scatter plot shows UMAP results of single cells based on scDaPars recovered APA profiles. Cells are labeled based on cell differentiation time points given in the original publication. (b) Cell-by-cell similarities represented by similarity matrices generated by R package SNFtool. (c) Scatter plots of UMAP results of cells in 96h of differentiation based on scDaPars results (left) and imputed gene expression (right). Cells are labeled by results from (b). (d) Scatter plot shows mean PDUI values of genes in subpopulation 2 (x-axis) and subpopulation 1 (y-axis). Genes with significant 3’UTR shortening and lengthening (PDUI differences >= 0.2) in subpopulation 2 are labeled in blue and red respectively. Bar plot shows the number of genes with significant shortening or lengthening in subpopulation 2 and p-value is calculated using single-tailed binomial test. (e) Selected GO terms enriched in the upregulated genes in subpopulation 2. (f) Example gene expression levels in two subpopulations. (g) Stream plot from STREAM which shows cell density along different trajectories at a given pseudotime.

Next, we investigated whether APA could help delineate novel cell subpopulations invisible to gene expression analysis alone. Imputation based on observed gene expression has been shown to enhance the identification of cell subpopulations (Li and Li 2018). Therefore, to ensure APA is providing additional information beyond expression, we first recovered plausible single-cell gene expression data using scImpute (Li and Li 2018), a state-of-the-art gene expression imputation method. Notably, although the imputed gene expression profile outputs more compact clusters than the raw expression, single cells collected from 72 and 96 h of differentiation were still largely overlapped (Supplementary Fig. 8). To characterize additional cellular heterogeneity, we integrated APA information with imputed gene expression using similarity network fusion (SNF) (Wang et al. 2014). By creating and converging separate similarity networks for APA dynamics and gene expression, SNF reduced noisy inter-cluster similarities among cells in 12 and 24 h of differentiation and enhanced intra-cluster similarities observed in one or both similarity networks (Fig. 6b). We then quantitatively compared the clustering results by using spectral clustering algorithm (Ng et al. 2002) on different similarity networks with the number of clusters *k* = 6. The clustering results are evaluated by normalized mutual information (NMI) (Witten et al. 2016) where *NMI* = 1 indicates a perfect match between the clustering results and the known differentiation time points. While gene expression imputation increased NMI from 0.76 to 0.85, integration of APA dynamics with imputed gene expression further increased NMI from 0.85 to 0.89, suggesting the benefits of adding APA information.

Besides unifying the clustering results of APA and gene expression, the fused similarity network also revealed novel and potentially meaningful subpopulations. For example, cells at 96 h of differentiation were divided into two previously unidentified subpopulations (Fig. 6b). Through analyzing APA and gene expression between the two subpopulations, we found that subpopulation 2, which was more distinct from cells in 72 h of differentiation than subpopulation 1, exhibited global 3’UTR lengthening compared to subpopulation 1 (P = 3.64e-8) (Fig. 6c, d); whereas the gene expression profile failed to distinguish the two subpopulations (Fig. 6c). APA profile quantified by bulk DaPars also failed to identify the 2 subgroups (Supplementary Fig. 9), indicating the superiority of scDaPars.

Since subpopulation 2 showed global 3’UTR lengthening, we hypothesized it may represent a more differentiated cell subgroup. To test our hypothesis, we performed differential gene expression analysis between subpopulation 1 and 2 using DESeq2 (Love et al. 2014). As a result, subpopulation 2 was characterized by higher expression of endoderm development marker genes including GATA6, EOMES, and SOX17 (Chu et al. 2016) (Fig. 6f, Supplementary Table 3). In addition, the transcriptional profile of subpopulation 2 also included significantly upregulated endoderm development related genes like LHX1, which is important for renal development (Reidy and Rosenblum 2009), and HGMA2, which is required for epithelium differentiation during embryonic lung development (Singh et al. 2014), suggesting it has a more differentiated phenotype than subpopulation 1. To further elucidate the global biological differences between the two subpopulations, we performed gene oncology (GO) analysis (Luo et al. 2009). We found that several endoderm development related GO terms were highly enriched in the upregulated genes in subpopulation 2 (Fig. 6e). Furthermore, using the expression of differential APA genes, we were able to separate the 2 subpopulations (Supplementary Fig. 10), indicating that some biologically meaningful subpopulations were masked by overall gene expression analysis. Finally, we conducted a trajectory analysis by STREAM (Chen et al. 2019) to independently show the validity of the identified subpopulations. Using cells at 0 h of differentiation as a natural starting point (root), we found that most cells are projected onto the inferred branches according to their corresponding differentiation time points (Supplementary Fig. 11a, b), and the derived pseudotime progression corroborated that cells in subpopulation 2 are more differentiated than those in subpopulation 1 (Fig. 6g, Supplementary Fig. 11c). Considered collectively, scDaPars calculated APA usage offered an additional layer of information in characterizing cellular heterogeneity that was otherwise invisible in gene expression analysis.

## Discussion

Here, we developed scDaPars, a novel bioinformatics algorithm to *de novo* identify and quantify single-cell dynamic APA events using standard scRNA-seq data. Many methods have been developed to measure the relative APA usages in RNA-seq data from bulk samples (Xia et al. 2014). However, the widespread dropout events in scRNA-seq data impede these bulk-sample based methods to quantify APA dynamics among single cells (Figs. 2d and 2e). To address this technical challenge in scRNA-seq, scDaPars first quantifies raw APA dynamics based on the two-poly(A)-site model introduced in DaPars (Xia et al. 2014). Since APA exhibits a cell-type specific pattern (Velten et al. 2015; Kim et al. 2019), scDaPars then clusters cells into different cell neighbors based on their calculated raw APA profiles. Next, scDaPars imputes missing APA usage by borrowing APA information of the same gene from neighboring cells. Benchmarking on both real and simulated data show the accuracy of scDaPars in predicting poly(A) sites, the ability in recovering missing APA usages, and the robustness in identifying dynamic APA events across cell types (Fig. 2 and 3).

Previously, methods for analyzing APA dynamics using scRNA-seq data mostly address the high technical noise in scRNA-seq data by creating pseudo-bulk RNA-seq data (i.e. pooled reads from cells that are assigned to the same cell cluster) (Shulman and Elkon 2019; Ye et al. 2020). Unlike scDaPars, even though these methods perform on scRNA-seq data, they do not quantify APA dynamics at the single-cell resolution but rather measure cell-cluster APA dynamics, which contradicts the purpose of single-cell sequencing (Supplementary Table 1). Additionally, previous methods are confined by cell cluster assignments determined by conventional gene expression analysis. In contrast, scDaPars quantifies single-cell APA dynamics independent of gene expression, which provides an additional layer of APA information that helps identify hidden cell states (Fig. 6c).

Finally, unlike existing methods, we expect scDaPars to be widely applicable to any scRNA-seq datasets. While the main analysis presented in this paper builds on scRNA-seq data generated by low-throughput SMART-Seq2 (Picelli et al. 2013) protocol, scDaPars can also be applied to large datasets generated by high-throughput droplet-based methods, e.g. 10X Chromium (Zheng et al. 2017). For example, scDaPars successfully revealed cell-type specific APA patterns in 3362 PBMCs sequenced by 10X Chromium (Ding et al. 2020) (Fig. 4a). Together, scDaPars provides an additional layer of APA information that helps identify cell subpopulations invisible to conventional gene expression analysis.

## Methods

### *De novo* quantification of dynamic APA events

scDaPars first performs *de novo* identification and quantification of dynamic APA events based on the two-poly(A)-site model introduced in DaPars. The bedgraph files for each single cell were used as input and jointly analyzed to calculate the APA dynamics measured as the Percentage of Distal PAS Usage Index (PDUI). For each gene, the distal PAS was identified as the end point of the longest 3’ UTR among all scRNA-seq samples, and the proximal PAS was inferred by optimizing the following linear regression model:

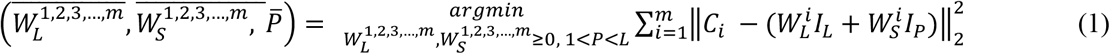

where 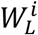 and 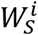 are the abundances of transcripts with distal and proximal PAS for cell *i, C_i_* is the read coverage of cell *i* normalized by total sequencing depth, *L* is the length of the longest 3’UTR, *P* is the length of the alternative proximal 3’UTR to be inferred, *I_L_* and *I_P_* are two indicator functions for long and short 3’UTRs such that 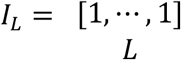 and 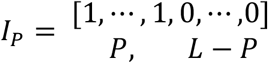. The optimal proximal PAS is selected by minimizing the deviation between the observed read density *C_i_* and the expected read density 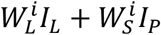 in all single cells. The APA dynamics are then quantified as PDUI for each gene in each single cell, with PDUI defined as:

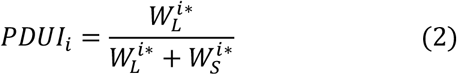

where 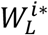 and 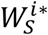 are the optimal expression levels of transcripts with distal and proximal PAS for cell *i*. The smaller the PDUI is, the less distal PAS is used, the shorter the 3’UTRs. The final output is a PDUI matrix in which rows represent genes and columns represent cells. Additionally, PDUIs can only be calculated in this step for genes with sufficient read coverage, which automatically separate genes into robust genes and dropout genes for future references.

### Detection of potential neighboring cells and outliers

Since APA exhibits alterations in different cell types and cell states in a global scale, scDaPars recovers missing single-cell level APA dynamics by borrowing information of the same gene from neighboring cells. A critical step here is to determine which cells are from the same cell subpopulation and therefore are neighboring cells. However, due to the technical limitation of scRNA-seq data, it is unlikely to completely cluster cells into true subpopulations based on the sparse PDUI matrix generated in last step. Instead, the goal of this step is to determine a set of potential neighboring cells which scDaPars will fine-tune in the following imputation step.

To increase the robustness and reliability of the clustering results and to find more plausible neighboring cells, scDaPars applies principle component analysis (PCA) to the raw PDUI matrix. While the PDUI matrix is sparse, the modularity of dynamic APA provides redundancy in gene dimensions, which can be exploited. Therefore, scDaPars selects principal components (PCs) that can together explain at least 40% of the variance in the data. Note that the neighboring cells are identified in these PCA dimensions while the imputation is performed on the full PDUI matrix.

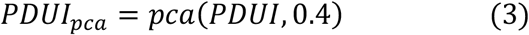

Next, scDaPars identifies and removes outlier cells from the analysis. The outlier cells may be the result of technical errors or may represent true rare biological variations, in either case, scDaPars will not use these outlier cells to impute missing APA dynamics in other cells. We calculate the distance matrix *D_N×N_* between cells based on the PCA transformed data *PDUI_pca_*. For each cell *m*, we define the Euclidean distance of cell *m* to its nearest neighbor as *d_m_*, resulting a set ***d*** = {*d*_1_,⋯, *d_N_*}. We denote the first quantile of ***d*** as *Q*_1_ and its third quantile as *Q*_3_ and the distance between *Q*_1_ and *Q*_3_ as interquartile range *IQR*. The outlier cells are defined as cells which are separated by more than 1.5 *IQR* to the third quantile *Q*_3_.

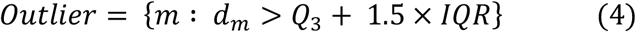

The remaining non-outlier cells {1,⋯, *N*}\*Outlier* are then clustered into subpopulations using graph-based community detection algorithm. The single cells are the vertices in the graph, and community detection in graphs will identify groups of vertices with high probability of being connected to each other than to members of other groups. We use R package *RANN* with default parameters to first identify the approximate nearest neighbors and convert neighbor relation matrix into an adjacency matrix. We then use *igraph* (*Csardi and Nepusz 2006*) to represent the resulting adjacency matrix as a graph and apply *walkstrap* (*Pons and Latapy 2005*) *algorithm* to identify communities of vertices (cells) that are densely connected. Suppose scDaPars divides cells into *K* subpopulations in this step, for each cell *m*, its potential neighboring cells *N_m_* are the other cells in the same cell subpopulation *k*.

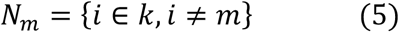

### Imputation of missing APA dynamics

After potential neighboring cells *N_m_* for each cell are determined, we impute APA dynamics cell by cell. Recall that PDUIs can only be estimated for genes with sufficient read coverage, scDaPars thereby automatically separates genes into robust genes and dropout genes when calculating the PDUI matrix. Here, we denote the set of robust genes for cell *m* as *R_m_* and the set of dropout genes that will be imputed in this step as *D_m_*. scDaPars then learns the cells’ similarities through the robust gene set *G_Robust,m_* and impute the APA dynamics of *D_m_* by borrowing information from the same gene’s APA dynamics in other neighboring cells learned from *R_m_*. To fine-tune the grouping of neighboring cells from *N_m_*, we use non-negative least squares (NNLS) regression:

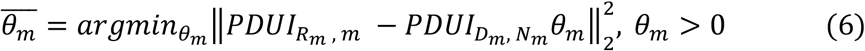

where *N_m_* represents the indices of cells that are potential neighboring cells of cell *m, PDUI_Gener_robust, m__* is a vector of response variables representing *R_m_* rows in the *m*-th column (cell *m*) of the original PDUI matrix, *PDUI_D_m_, N_m__* is a sub-matrix of the original PDUI matrix with dimensions |*D_m_*| × |*N_m_*|. The goal is to find the optimal coefficients 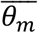 of length |*N_m_*| that can minimize the deviation between APA dynamics of *R_m_* in cell *m* and those in potential neighboring cells. The advantage of using NNLS is that it has the property of leading to a sparse estimate of *θ_m_*, whose components may have exact zeros, so that true neighboring cells of cell *m* are conveniently selected from *N_m_*. Once 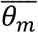 is computed, we have a vector of weighted neighbors associated with each cell in our data. scDaPars use this coefficient 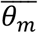 estimated from the set *R_m_* to impute the APA dynamics of genes in the set *D_m_* in cell *m*, thus coupling the tasks of borrowing APA information of the same gene in neighboring cells and of transferring APA information from robust genes *R_m_* to dropout genes *D_m_*.

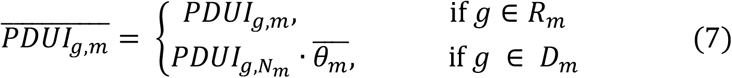

### Preprocessing of scRNA-seq data

All the scRNA-seq datasets used in this study are summarized in Supplementary Table 4. For low-throughput datasets generated by SMART-Seq2 (Picelli et al. 2013) protocol, we downloaded the publicly available fastq files from GEO database and aligned the reads using STAR 2.5.2 (Dobin et al. 2013) with default parameters, generating one BAM file for each single cell. For high-throughput datasets generated by 10X Chromium (Zheng et al. 2017), we downloaded the fastq files and aligned the reads using Cell Ranger 3.0.2. We then selected reads with correct unique molecular identifier (UMI) using Drop-seq tools *FilterBAM* (*Macosko et al. 2015*) and remove reads with duplicated UMIs using UMI-tools *dedup* (Smith et al. 2017). We next merged reads originated from same cells together and generated one BAM file for each single cell. The BAM files are used as inputs for subsequent scDaPars analysis.

### Generation of synthetic dataset

The synthetic dataset was created based on bulk RNA-seq data generated from 13 immune cell types (Schmiedel et al. 2018). The different immune cell types are isolated so that each sample only contains cells from one cell type. We used DaPars to estimate the APA usage in these bulk samples and generated an APA matrix, in which rows represent genes and columns represent samples. Since widespread dynamic APA events were reported in Naïve and activated CD4 T cells, we selected only samples that belong to these two cell types for the following simulation.

We down-sampled the resulting bulk APA matrix to emulate the APA profiles generated from single-cell data. We first calculated the dropout rate for each gene in the benchmark immune dataset (Ding et al. 2020). Next, for each gene in the bulk APA matrix, the dropout rate is randomly selected from the set of real dropout rates with replacement. Finally, we used Bernoulli distribution with p equals to the selected dropout rate and n equals to the number of samples to introduce dropouts into the synthetic dataset. The final dropout introduced data has a ~50% dropout rate. Notice that the generation of the synthetic dataset is independent from the models of scDaPars, so that it can be used to evaluate scDaPars in a fair way.

### Benchmark comparison of scDaPars

To illustrate the advantage of scDaPars, we applied scDaPars, scAPA and Sierra to a benchmark 10X Chromium dataset containing 3362 PBMCs (Ding et al. 2020). scAPA measures differential usage of poly(A) sites between different cell types by the proximal poly(A) site usage index (proximal PUI). Since we want to test scAPA’s ability for quantifying single-cell-level APA usage, we input single-cell coverage into scAPA to generate a cell by transcript proximal PUIs matrix to perform the clustering analysis. The Sierra pipeline does not yield PDUI like measurements. Instead, it generates a peak count matrix in which peak coordinates are annotated according to the genomic features they fall on including UTRs, exons, or introns. In order to calculate APA usage from the peak count matrix, we first selected peaks falling on the 3’UTRs and only kept transcripts with more than one peak. We then transferred the peak count matrix into an APA matrix by calculating the relative usage of the most distal peak. The resulting APA matrix were used for the clustering analysis. Finally, we performed silhouette analysis by *silhouette()* in R package *cluster* v2.1.0. to quantitatively evaluate the clustering accuracy of the three methods.

## Data access

The scRNA-seq datasets used in this manuscript are all publicly available and are summarized in Supplementary Table 4. The 2 single-cell PBMC data are available at the Gene Expression Omnibus (GEO) under accession code GSE132044. The breast cancer data are available at GEO under accession code GSE75688. The time-course definitive endoderm data are available at GEO under accession code GSE75748. The DICE immune data used to generate synthetic dataset were obtained from dbGaP under study accession code phs001703.v1.p1. The R package scDaPars is freely available at https://github.com/YiPeng-Gao/scDaPars.

## Author contributions

W. L. conceived and supervised the project. Y.G. performed the data analysis. Y.G., L.L., W.L. interpreted the data. Y.G., L.L., W.L., C.A. wrote the manuscript.

## Acknowledgements

We thank Yikai Luo, Dr. Joel Neilson at Baylor College of Medicine, members of the Li lab at University of California, Irvine, and Dr. Jingyi Jessica Li at University of California, Los Angeles for insightful discussions. We thank Dr. Chen Chao at Baylor College of Medicine for his suggestions on this manuscript. This work is supported by US National Institutes of Health (NIH) grants R01HG007538, R01CA193466 and R01CA228140 to W.L. and the Cancer Prevention Research Institute of Texas (CPRIT) grant RR170048 to C.A. C.A. is a CPRIT research scholar.

## Supplementary Tables

**Supplementary Table 1.** Comparison of different scRNA-seq APA quantification methods

| Method | Single-cell level | Single-gene level | Cell-type-label free |
| --- | --- | --- | --- |
| scDaPars | ✓ | ✓ | ✓ |
| scDAPA |  | ✓ |  |
| scAPA |  | ✓ |  |
| Chung et al. | ✓ |  | ✓ |

**Supplementary Table 2.** Differential gene expression summary table for B cell proliferation signature genes between group 1 and group 2 B cells.

| Gene Symbol | Log2 Fold Change | P-value | Gene Symbol | Log2 Fold Change | P-value |
| --- | --- | --- | --- | --- | --- |
| CCNE2 | 8.637907005 | 4.665E-09 | PLK4 | 2.243609617 | 2.724E-02 |
| CCNA2 | 4.803637943 | 9.399E-07 | GTSE1 | 2.033800133 | 2.849E-02 |
| NEK2 | 5.043927629 | 9.463E-07 | MCM3 | 2.041564617 | 4.137E-02 |
| CDC20 | 4.495114905 | 1.630E-06 | AURKB | 1.870984406 | 4.462E-02 |
| MCM4 | 3.991918849 | 2.174E-06 | CCNB1 | 1.660149707 | 5.296E-02 |
| CCNF | 4.105056458 | 4.720E-06 | AURKA | 2.024196574 | 7.011E-02 |
| MCM6 | 3.806024739 | 6.365E-06 | BUB1B | 1.690663297 | 8.141E-02 |
| KIF11 | 3.305184966 | 8.099E-05 | GADD45A | 2.498681071 | 8.849E-02 |
| CDK5 | 5.729766087 | 9.758E-05 | CENPF | 1.397881644 | 1.135E-01 |
| CDC45 | 3.846374532 | 3.668E-04 | CENPE | 1.233362375 | 1.252E-01 |
| CDK1 | 2.862335992 | 8.109E-04 | TFDP1 | 1.387714487 | 1.271E-01 |
| TTK | 3.664301687 | 1.292E-03 | CCNB2 | 1.33876417 | 1.293E-01 |
| KIF22 | 2.40064932 | 2.060E-03 | RPA3 | 0.942772863 | 2.137E-01 |
| CENPA | 4.606155804 | 2.292E-03 | KIF23 | 1.253952495 | 2.247E-01 |
| MKI67 | 2.33120824 | 2.360E-03 | CDC6 | 1.323104442 | 2.416E-01 |
| CHEK1 | 2.61335517 | 4.228E-03 | CDC25B | 1.138407673 | 3.378E-01 |
| TPX2 | 2.406900695 | 5.406E-03 | PTTG1 | 0.637142701 | 3.651E-01 |
| RFC3 | 2.785858262 | 5.984E-03 | MAD2L1 | 0.618252654 | 4.851E-01 |
| TMPO | 1.964265986 | 8.086E-03 | WEE1 | 0.402150684 | 5.184E-01 |
| PCNA | 2.235343755 | 8.233E-03 | E2F5 | 0.530712699 | 6.110E-01 |
| UBE2C | 2.331979017 | 9.358E-03 | RAD17 | -0.077271021 | 9.172E-01 |
| CDKN2C | 3.765539392 | 1.057E-02 | GADD45B | -0.060809059 | 9.384E-01 |
| ZW10 | 5.01976692 | 1.350E-02 | RABGAP1 | 0.023630566 | 9.796E-01 |

**Supplementary Table 3.**
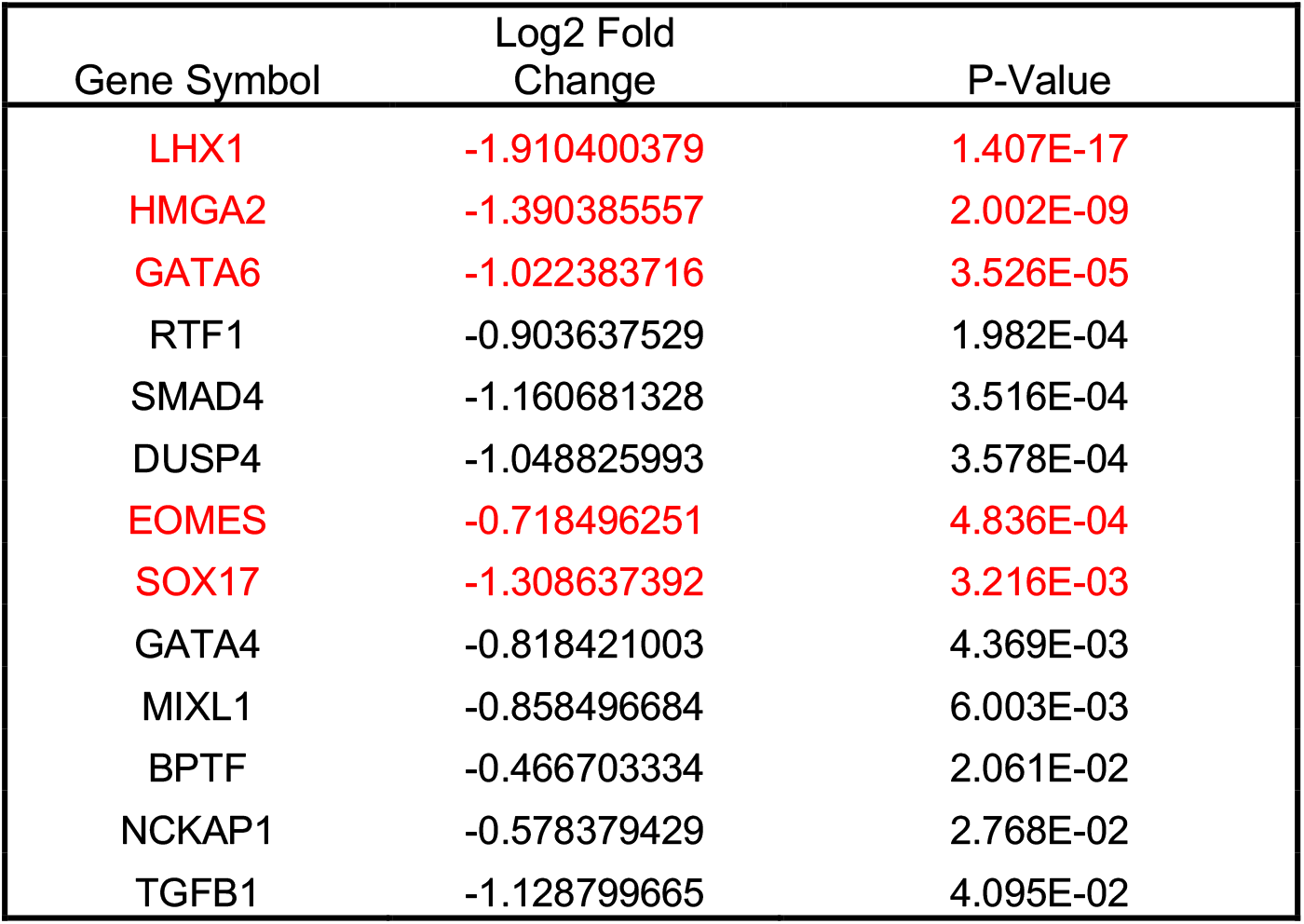
Significantly upregulated genes in subpopulation 2 that are annotated with the GO term “endoderm development.”

| Gene Symbol | Log2 Fold Change | P-Value |
| --- | --- | --- |
| LHX1 | -1.910400379 | 1.407E-17 |
| HMGA2 | -1.390385557 | 2.002E-09 |
| GATA6 | -1.022383716 | 3.526E-05 |
| RTF1 | -0.903637529 | 1.982E-04 |
| SMAD4 | -1.160681328 | 3.516E-04 |
| DUSP4 | -1.048825993 | 3.578E-04 |
| EOMES | -0.718496251 | 4.836E-04 |
| SOX17 | -1.308637392 | 3.216E-03 |
| GATA4 | -0.818421003 | 4.369E-03 |
| MIXL1 | -0.858496684 | 6.003E-03 |
| BPTF | -0.466703334 | 2.061E-02 |
| NCKAP1 | -0.578379429 | 2.768E-02 |
| TGFB1 | -1.128799665 | 4.095E-02 |

**Supplementary Table 4.** Summary of scRNA-seq data used in this manuscript

| Data | Sequencing Protocol | # of Single cells | GEO accession code |
| --- | --- | --- | --- |
| PBMC1 | SMART-seq2 | 384 | GSE132044 |
| Breast Cancer | SMART-seq2 | 563 | GSE75688 |
| Definitive Endoderm | SMART-seq2 | 758 | GSE75748 |
| PBMC2 | 10X Chromium | 3362 | GSE132044 |

